# Activated CD8 T cells express two distinct P-Selectin Ligands

**DOI:** 10.1101/167957

**Authors:** Douglas A. Carlow, Michelle C. Tra, Hermann J. Ziltener

## Abstract

**One sentence summary:** Murine primary in-vivo activated CD8+ T cells express two ligands for P-selectin, canonical PSGL-1 and a cell-extrinsic ligand docked on L-selectin.

**Abstract:** P-selectin (PSel) expressed on activated endothelia and platelets supports recruitment of leukocytes expressing PSel ligand (PSelL) to sites of inflammation. While monitoring PSelL expression on activated CD8^+^ T cells (Tact) in adoptive transfer models, we observed two distinct PSelL on responding donor cells, the canonical cell-intrinsic PSelL PSGL1 and a second undocumented PSelL provisionally named PSL2. PSL2 is unusual among selectin ligands in that it is cell-extrinsic, loaded onto L-selectin (LSel) expressed by Tact but not LSel on resting naïve CD8^+^ T cells. PSL2 expression is highest on Tact responding in peripheral lymph nodes and low on Tact responding in spleen suggesting that the original source of PSL2 is high endothelial venules, cells known to produce LSelL. When both PSGL1 and PSL2 were absent from the surface of Tact, no significant residual PSelL activity was detected. PSL2 is a ligand for both PSel and LSel and can physically bridge the two selectins. The LSel/PSL2 complex can mediate PSel-dependent adherence of Tact to immobilized PSel-hIgG or to activated platelets, either independently or cooperatively with PSGL1. PSel engagement of PSGL1 and LSel/PSL2 would likely deliver distinct signals known to be relevant in leukocyte recruitment.

## Introduction

Leukocyte tethering to endothelium is the initial step in movement of leukocytes from blood into tissue, a fundamental process in lymphoid homeostasis, the inflammatory response, and immunological defense. These tethering interactions begin with low affinity contacts between leukocytes and activated vascular endothelia through binding of selectins to their ligands on opposing cell surfaces. Identification of all physiologically relevant ligands is needed to complete the understanding of selectin function in the aforementioned fundamental processes.

P-Selectin (PSel) and E-selectin (ESel) *(1)* are expressed on activated endothelium and tether to ligands expressed on leukocytes to support their recruitment during inflammation *(2-4)*. PSel is also expressed at high density on activated platelets and cyclically on thymic endothelium paralleling thymic receptivity in homeostasis *(5)*. All selectins recognize ligands modified with sLex tetrasacharides but engage largely distinct ligand sets determined by additional modifications of the sLex glycan and properties of the scaffold or peptide backbone. PSel is generally thought to have a single, broadly utilized and physiologically active ligand, Platelet Selectin Glycoprotein Ligand 1 (PSGL-1). PSel recognition of PSGL1 dimer requires sLex modified branched O-glycans generated in the golgi by the core 2 C2GnT1 enzyme, together with sulfated tyrosine residues adjacent to the O-glycan attachment site. Such cell intrinsic ‘decorated’ PSGL1 P-selectin ligand (PSelL) is present constitutively on neutrophils but induced on T lymphocytes only after their antigen-driven activation in secondary lymphoid organs. Induction of PSGLl-PSelL expression constitutes part of the cell-intrinsic response by lymphocytes to support recruitment via PSel on vasculature of inflamed tissue.

L-selectin (LSel) supports steady-state lymphocyte homing to peripheral lymph nodes through multiple glycoprotein ligands (LSelL) expressed on high endothelial venules (HEV). LSelL on HEV are modified with directly sulfated sLex glycans (6-O-sulfated sLex)(6-9) presented on extended core 1 O-glycans, C2GnT1 branched O-glycans(10), or N-glycans *(11, 12)*. Glycosyltransferases and sulfotransferases restricted to HEV (7-9) generate the 6-O-sulfated sLex typically required for recognition by LSel. Apart from its role in LN homeostasis, LSel is also active in chronic inflammatory contexts where endothelium of inflamed tissue can differentiate to an HEV-like phenotype expressing LSelL that drives heavy leukocyte recruitment and formation of tertiary lymphoid structures *(13)*. LSel’s contribution in primary acute inflammatory recruitment prior to such endothelial ‘conversion’ is not widely recognized because LSelL are not detected on tissue vascular endothelium. In these circumstances, LSel can tether to decorated PSGL1 on leukocytes or leukocyte fragments previously adhered to endothelium in a process called secondary capture (14).

While studying formation of PSGL1-PSelL on primary in vivo activated CD8^+^ T cells (Tact) we detected a PSGL1-independent PSelL we provisionally named P-Selectin-ligand-2 (PSL2). As analysis of PSL2 on Tact progressed, aspects of its expression were increasingly consistent with it participating in recruitment of this major lymphocyte subset. T cell immunity directed at peripheral sites is generated in PerLN and this process is normally accompanied by induction of the canonical leukocyte-expressed PSelL PSGL1. Like PSGL1, PSL2 was reliably detected on CD8^+^ T cells after they were activated by antigen in peripheral lymph nodes (PerLN). The contemporaneous expression of both decorated PSGL1 and PSL2 PSelL on Tact positions them to participate/cooperate in processes supported by PSel during recruitment.

## Results

### Discovery of PSL2

The current paradigm for induction of P-selectin ligand on CD8^+^ T cells begins with their antigen-driven activation in secondary lymphoid organs and, in the absence of additional signals such as retinoic acid in the gut *(15, 16)*, the subsequent induction of glycosyl transferases including C2GnT1 that synthesize sLex-modified Core 2 branched O-glycans on PSGL1 required for recognition by PSel *(17,18)*, as shown in *Figure 1A*. This decorated PSGL1 serves as the canonical ligand for PSel the latter being expressed on both activated platelets and activated endothelium in inflammatory processes. When CD8^+^ T cells are activated by dendritic cells in vitro using two independent T cell receptor transgenic models, OT1 responses specific for ovalbumin and HY responses specific for male antigen, this pattern of PSGL1-PSelL formation is recapitulated precisely. As shown in *Figure 1B*, formation of PSelL on *in vitro* activated CD8^+^ T cells was strictly dependent on T cells expressing both PSGL1 and C2GnT1, consistent with PSGL-PSelL being the sole PSelL. However, on primary *in vivo* activated CD8^+^ T cells (Tact) PSel-hIgG staining persisted on responding T cells deficient in either PSGL1 or C2GnT1 but was lost when EDTA was included in the staining procedure_ suggesting that another PSelL was present on in vivo generated Tact; this ligand was given the provisional name PSL2.

**Figure 1:**
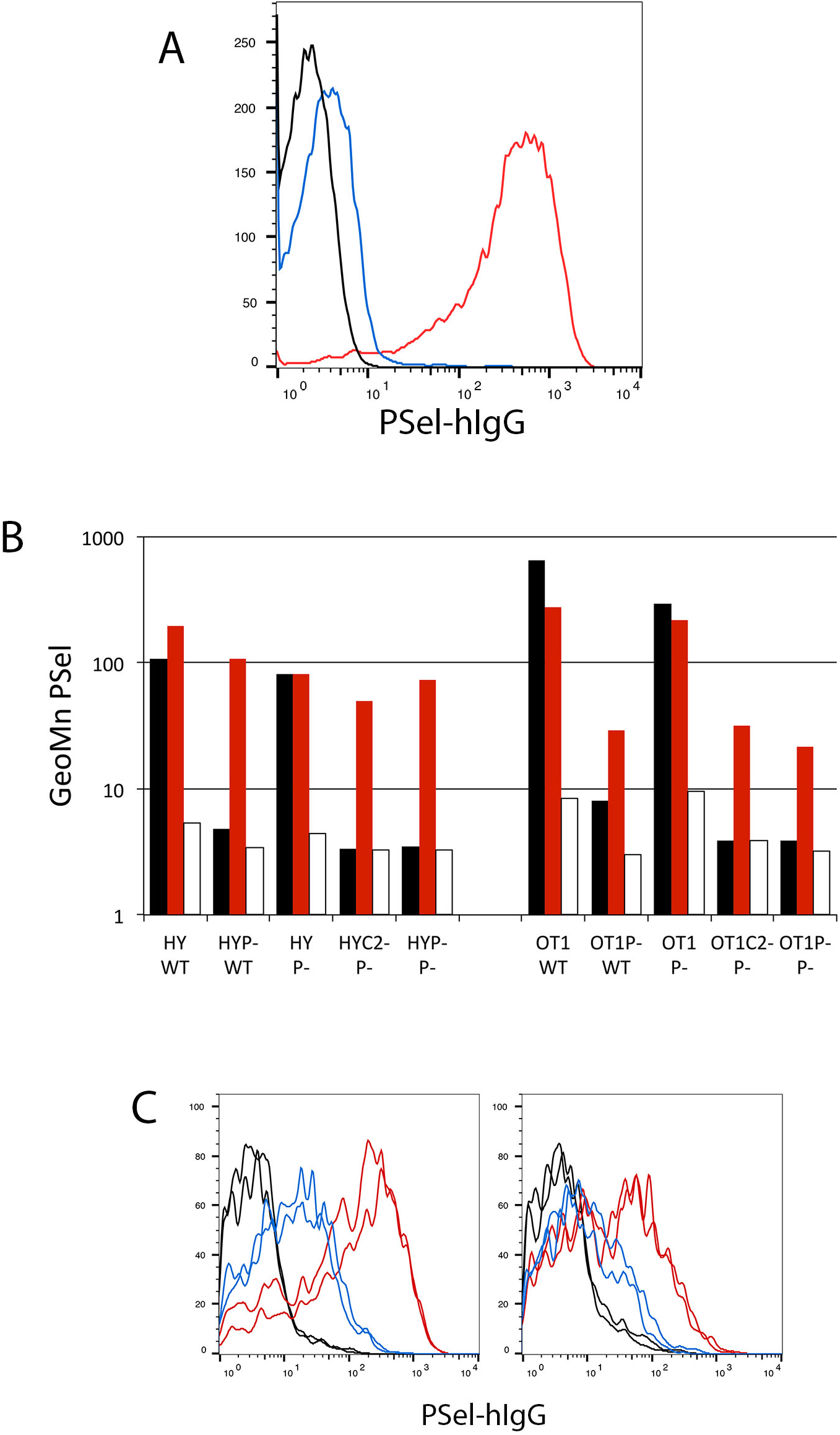
Detection of a PSGL1-independent PSelL on in-vivo activated T cells. **(A) Formation of PSGL1-PSelL requires C2GnT1:** Spleen cells from *B6* (red), *PSGL-1^nul1^* (black), or *C2GnT1^nul1^* (blue) mice were activated with Concanavalin A for two days, subcultured with fresh media and interleukin 2 for two more days, and stained with PSel-hIgG and anti-hIgG-PE. Cytometer gain was set such that fluorescence of unstained controls (not acquired) were comparable with fluorescence of *PSGL1^nul1^* cells. **(B) PSL2, a PSGL1-independent PSelL, is detected on CD8 T cells activated *in vivo*:** PSel-hIgG staining of day-3 activated CD8^+^ T cells from HY male antigen specific, or OT1 ovalbumin specific, T cell receptor transgenic mice responding to antigen *in vitro* or *in vivo*. CFSE-labeled responding cells on *wild type (B6=WT), PSGL-1^nul1^ (P-)*, or *C2GnT1^nul1^* (C2-) genetic backgrounds were stimulated *in vitro* with antigen presenting (male or Ova-pulsed) splenic dendritic cells (DCs) from B6 (WT) or *PSGL-1^nul1^* (P-) mice or adoptively transferred into B6 (WT) or *PSGL-1^nul1^* (P-) male recipients. After three days, responding cells were harvested from in vitro cultures *(black bars)* or in vivo responding donor cells in PerLNC from recipient mice *(red bars)*, and stained with PSel-hIgG chimera and anti-human IgG-PE. EDTA was included in parallel samples to confirm divalent cation (Ca^++^) dependent selectin binding *(open bars)*. **(C) PSL2 is preferentially expressed on Tact responding in peripheral lymph node.** PSel-hIgG staining of CFSE-labeled *HY-PSGL^null^* T cells responding to male antigen in *PSGL1^null^* male recipients *(left panel)* or CFSE-labeled *OT1-PSGL^null^* T cells responding to ovalbumin antigen in *PSGL1^null^* recipients *(right panel)*. Donor cells were recovered from peripheral lymph nodes (*red*) or spleen (*blue*) three days after adoptive transfer into respective recipients. Ex-vivo cells were subjected to gating for viable PI^neg^, CD8^+^, responding (CFSE-diluted) donor cells. Staining of duplicate recipient mice are shown for each donor. Control PSel-hIgG staining performed in the presence of EDTA shown for respective donor/recipient combinations *(black)*.

PSL2 was revealed on ex-vivo Tact generated from either *PSGL^nul1^* or *C2GnT1^null^* donor cells that are both unable to synthesize the PSGL1-PSelL for different reasons. HY and OT1 responses carrying these *null* alleles were used interchangeably in PSL2 analyses going forward although *C2GnT1^nul1^* donors have superior LN access and gave superior Tact yields at day 3 compared to *PSGL1^nul1^* donors (19). Finally, the use of *PSGL1^nul1^* mice *(or C2GnT1^null^mice)* as recipients/stimulators for activating antigen specific responses excluded the possibility of Tact acquisition of decorated PSGL1-PSelL from stimulating dendritic cells through trogocytosis. Some trogocytosis of PSGL1 can occur in these systems but was a minor contribution to PSelL detected on Tact (data not shown).

### PSL2 expression is predominant in PerLN but not spleen

PSL2 levels were compared on Tact harvested from PerLN and spleen in both OT1 and HY T cell receptor transgenic models as shown in *Figure 1C*. Surprisingly, the levels of PSL2 differed markedly on Tact from the two secondary lymphoid organs; responses in PerLN consistently exhibited higher levels of PSL2 than Tact responding in spleen. PSL2 levels on Tact responding in mesenteric lymph node (MesLN) were intermediate between PerLN and spleen (not shown).

### PSL2 expression is dependent on the presence of divalent cation

CD8^+^ T cells from *HY* mice (able to produce both PSGL1 and PSL2 PSelL) or *HY PSGL^nul1^* mice (able to produce only PSL2 PSelL) responded in male recipients by proliferation and expression of PSelL as shown in *Figure 2A*. A subset of CD8^+^ cells in the HY model express endogenous (non-transgenic) T cell receptor alpha chains, do not respond to male antigen, remain CFSE^high^, and do not express PSelL; these non-responding donor cells thus serve as a reference point for PSel-hIgG staining and proliferation of Tact. Expression of both PSGL1 and PSL2 PSelL required prior T cell activation. Surprisingly, brief exposure to EDTA-media at 4°C rendered Tact devoid of PSL2 expression even if Tact were returned to Ca^++^ replete conditions prior to PSel-hIgG staining as shown in *Figure 2A*. Pre-washing with calcium chelator EGTA also resulted in loss of PSL2 detection with PSel-hIgG (data not shown). This EDTA ‘pre-wash’ treatment was distinguished from controls where EDTA is included during PSel-hIgG staining to demonstrate calcium dependent binding by selectin. In the later case, EDTA prevents selectin binding to ligand by stripping Ca^++^ from the ligand-binding face of selectin, thereby disabling Ca^++^ ion mediated coordination between selectin and glycan atoms presented by ligand *(20)*. As shown in *Figure 2B* PSL2 expression was maintained relatively well when Tact were serially washed in minimal media with diminishing concentrations of Ca^++^ or Mn^++^ but was lost in media supplemented with Mg^++^.

**Figure 2:**
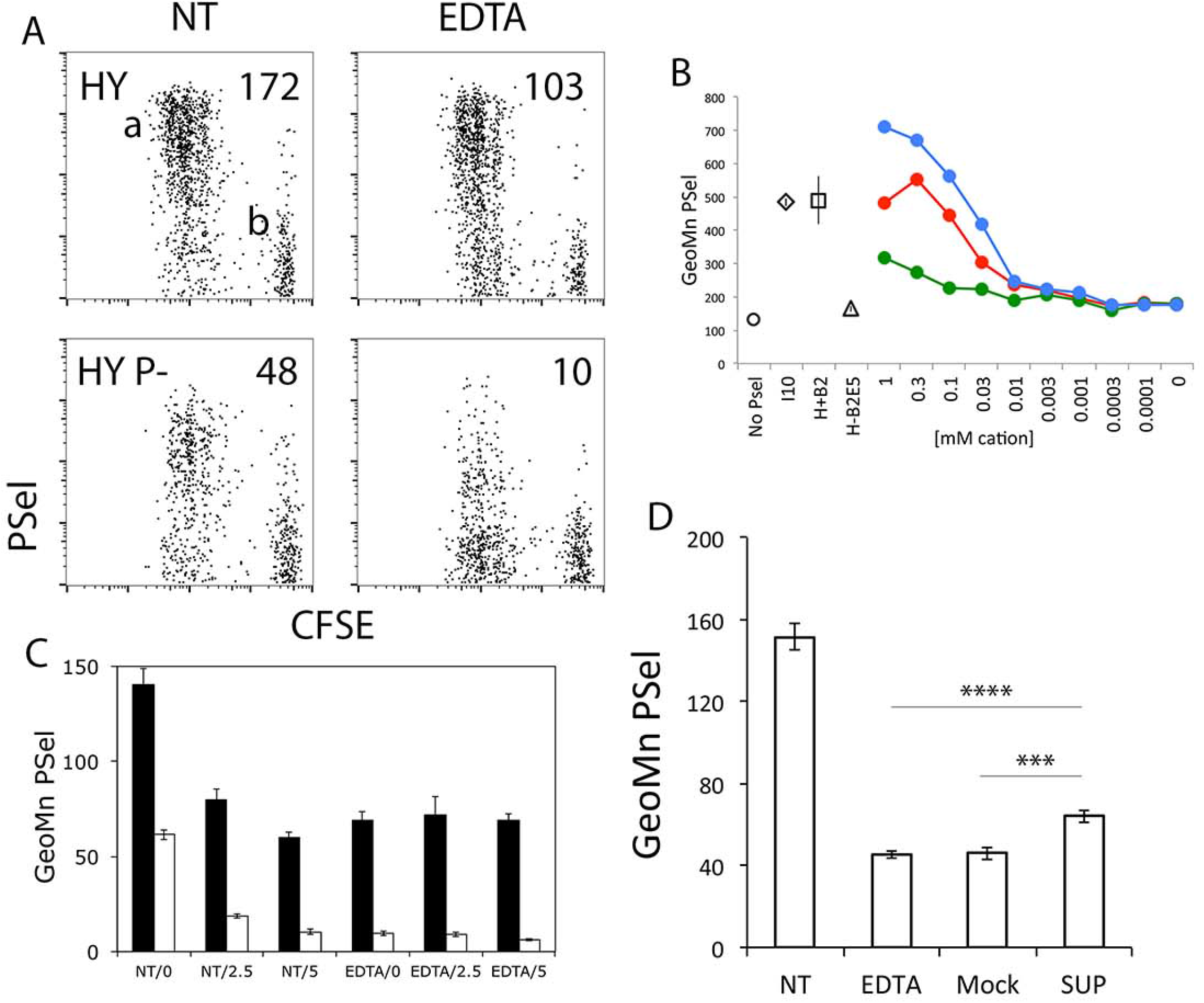
PSL2 association with the Tact cell surface requires Ca^++^. **(A) Cation removal eliminates expression of PSL2:** PSel-hIgG staining of ex-vivo CD8^+^ CFSE-labeled *HY* or *HY-PSGL1^null^* donor T cells responding (CFSE^low^ cloud ‘a’) and nonresponding (CFSE^high^ cloud ‘b’) at day 3 in *PSGL1^null^* recipients *(Ieft panels)*. In *right panels*, an aliquot of the same cells was washed twice in EDTA-containing media, returned to Ca^++^ replete media, and then stained with PSel-hIgG. Geometric mean values for PSel staining of responding (CFSE diluted) donor cells is shown. **(B) Cation specificity to support PSL2 expression:** PSL2CD8^+^ CFSE-labeled Tact from *OT1-C2GnT1^null^* responding in vivo to Ova were recovered at day 3 from *PSGL1^null^Thy1.1* recipients for PSel-hIgG staining on donor cells. The cation-specific dependence of PSL2 expression was assessed by 5x serial washes of cells in media with the indicated concentrations of cations, bIue-Mn^++^, red-Ca^++^, green-Mg^++^. *‘No PSel’* (control washed in I10 culture media and stained with all staining reagents except PSel-hIgG chimera), *I10* tissue culture media, *H+ B2* (Hanks+ with 2mg/ml BSA), *H-B2 E5* (Hanks^-^ with 2mg/ml BSA and 5mM EDTA). Standard deviation of triplicate stains shown. **(C) Persistence of PSGL1 PSelL and loss of PSL2 PSelL with *in vitro* culture:** HY (bIack coIumns) and *HY-C2GnT1^null^ (open coIumns)* donor Tact recovered from male *PSGL1^null^* recipients were either untreated *(NT)* or pre-washed with EDTA-media (PSL2 stripped). Cells were then transferred to I10 media and held on ice (0) or cultured at 37°C for 2.5 or 5 hours prior to PSel-hIgG chimera staining. Standard deviation of triplicate stains shown. **(D) PSL2 rebinding:** CD8^+^ *HY-C2GnT1^null^* donor Tact recovered from PerLN of male *PSGL1^null^ Thyl.1* recipients on day 3 were enriched (re: Mat. & Meth.) and either stained for PSel *(NT)* or treated with EDTA-media to generate donor cells lacking PSL2 *(EDTA)* and a supernate extract. This supernate was made Ca^++^ replete *(SUP)* and compared with a mock supernate, media prepared identically except never exposed to cells *(Mock)*, for capacity to restore PSelL on the Tact that had been pre-washed with EDTA-media. Mean and standard deviation of three PSel-hIgG staining replicates for each condition shown along with P values determined by two-tailed Student’s T-test. **** P value = 0.00073, *** P value = 0.0018.

The Ca^++^ dependence of PSL2 detection might reflect either (i) a cation requirement for a conformational domain on PSL2 used in PSel binding, or (ii) a cation requirement for PSL2 association with the cell surface. Ca^++^ could mediate PSL2 binding directly to cell surface lipids, as for Gal-domain containing proteins, or through an alternate docking molecule. When ex-vivo Tact were cultured briefly at 37°C, expression of PSL2 was lost while PSGL1 PSelL was maintained, *Figure 2C*. When Tact were pre-washed with EDTA-media and then cultured in Ca^++^ replete media, PSGL1 PSelL was maintained whereas PSL2 was lost in the wash and no recovery of PSL2 was observed. If PSL2 presented a conformational PSelL dependent on Ca^++^, recovery of PSL2 expression during culture might have been expected-but did not occur.

Alternatively, if treatment with EDTA eluted PSL2 from the cell surface into media, this eluted supernatant might restore PSL2 expression if ‘stripped’ Tact were subsequently reexposed to the supernatant in the presence of Ca^++^. As shown in *Figure 2D* this appeared to be the case as partial restoration of PSL2 expression relied on provision of both Ca^++^ and supernate. Based on these data we conclude that PSL2 is a PSelL bound through Ca^++^ to the surface of Tact. EDTA pre-washing strips PSL2 from Tact while leaving PSGL1 expression intact.

### PSL2 on Tact is cell-extrinsic and dependent on recipient C2GnT1 expression

Data in *Figure 1* indicated that PSL2 expression on in vivo generated Tact was independent of cell-intrinsic C2GnT1, in contrast to PSel recognition of PSGL1. Our initial interpretation of this difference was that the structure of the ligand on PSL2 must differ from the glycan array presented on PSGL1 for PSel recognition. However, when Tact were generated in recipients lacking *C2GnT1*, PSel detection of PSL2 was lost *(Figure 3A)*. This finding suggested that requirement of C2GnT1 for PSel recognition of PSL2 was indeed preserved and that PSL2 expressed on donor Tact was sourced from the recipient, and not a cell-intrinsic product of responding donor T cells. Sourcing of PSL2 from recipient tissue was also consistent with the observations that (i) T cells activated in vitro lack of PSL2 *(Figure 1)*, (ii) the association of PSL2 with Tact was Ca^++^ dependent and reversible, *(Figure 2A,B,D)*, (iii) PSL2 expression was rapidly and irreversibly lost during culture of ex-vivo Tact, a loss accelerated by brief washing in EDTA-media *(Figure 2C)*, and (iv) PSL2 expression on Tact was independent of cell-intrinsic *C2GnT1 (Figures 1A & 3A)*. Collectively these data demonstrated that PSL2 on Tact was cell-extrinsic and that the total PSelL generated on Tact during the primary response of OT1 CD8^+^ T cells in PerLN was the sum of the cell-intrinsic PSGL1 and cell-extrinsic PSL2; no additional ligand of PSel was visible in this system *(Figure 3A)*.

**Figure 3:**
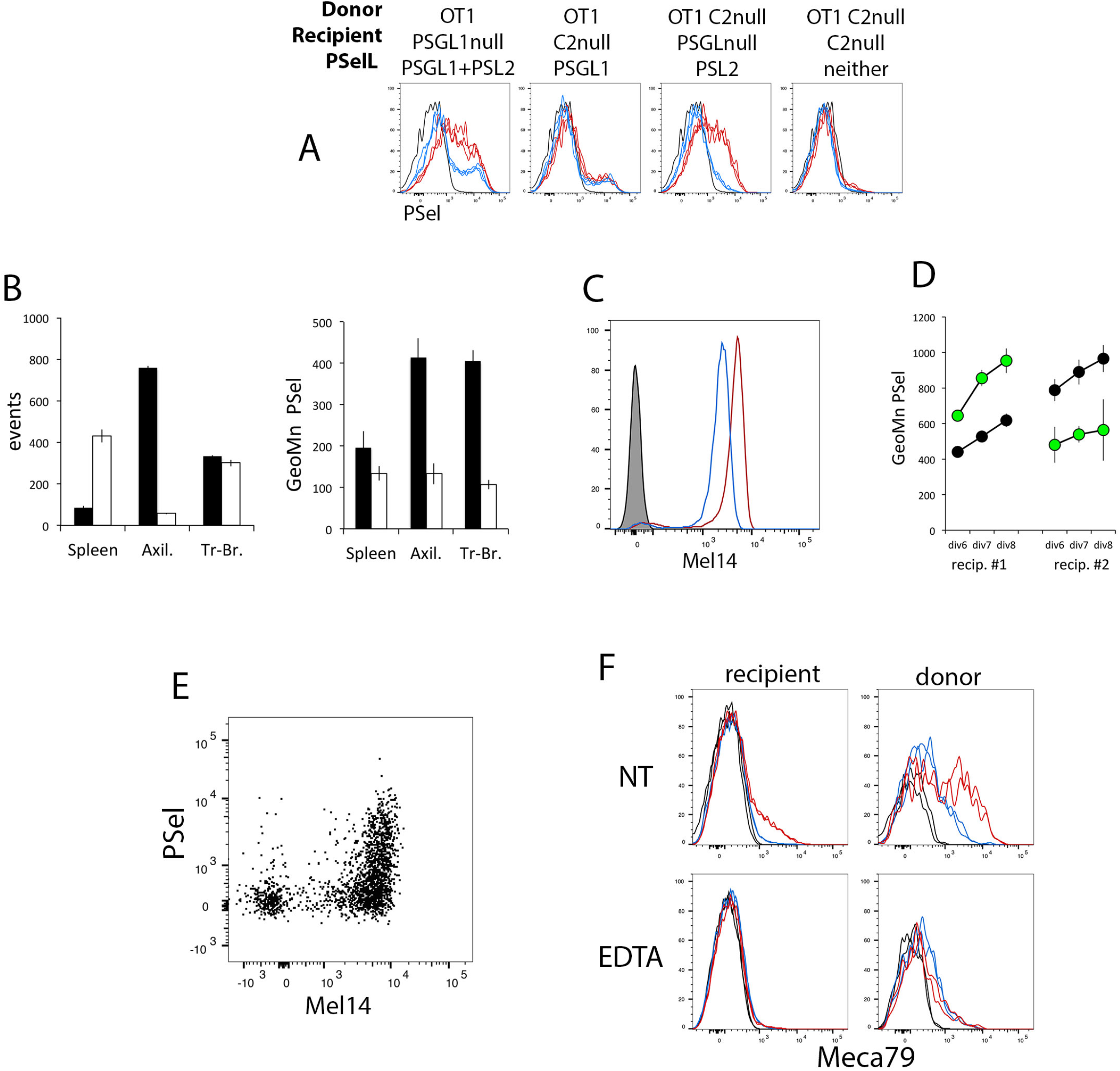
PSL2 is T cell-extrinsic, dependent on recipient C2GnT1 expression, and docks on L-selectin. **(A) PSL2 expression requires C2GnT1 expression in recipient and, together with PSGL1, constitute total PSelL on Tact:** CTV-labeled donor cells of genotypes indicated were injected into recipients with Ova antigen. Viable CD8^+^ responding donor cells were analyzed at day 3 for PSel-hIgG staining *(red*, untreated; *bIue*, after EDTA pre-wash; *bIack*, control stain without PSel-hIgG). **(B) LSel-independent entry of CD8 T cells into tracheo-bronchial LN:** OT1-PSGL^null^-LSeI^+/+^ and *OT1-PSGL-LSel’Z-* donor cells labeled with CTV or CFSE tracking dyes were co-injected iv onto *PSGL-Thyl.1* recipients that also received ova antigen. Spleen, axillary LNs, and tracheo-bronchial LNs were harvested at day 3 and assessed for relative numbers *(Ieft panel)* of *LSel^+/+^ (bIack coIumns)* and *LSel^-/-^ (open coIumns)* donor Tact, and PSelL expression *(rightpanel)*. Standard deviation of triplicate stains shown. **(C) LSel CD62L expression is gene dose dependent** Mel 14 staining of naïve LSeI^-/-^ (bIue Iine) and LSeI^+/+^ (red Iine) OT1 PSGL1^null^ **CD**8**+ T cells.** **(D) PSL2 expression on Tact responding in PerLN correlates with LSel gene dose:** Naïve *OT1-PSGL LSel^+/+^* and *OT1-PSGL1 LSel^-/-^* donor cells were labeled with CFSE or CTV, combined, and co-injected iv together with antigen (Ova) into *PSGL^nul1^ Thy1.1* recipients. PSel-hIgG staining shown at comparable cell division numbers for donor cells obtained from the same recipient at day 3. *Recipient #1, OT1-PSGL LSel+/+ (CFSE, green)* and *OT1-PSGL1 LSel^-/-^ (CTV, black). Recipient #2*, same donor cells but dyes reversed. Standard deviation of triplicate stains shown. **(E) Mel14 and PSel-hIgG co-staining of Tact:** *OT1C2GnT1^null^* donor cells in *PSGL1^null^* recipients responding at day 3 to Ova antigen in PerLN; analysis gated on responding (CFSE-diluted), viable, CD8^+^ donor cells. Standard deviation of triplicate stains shown. **(F) Meca 79 staining of CD8^+^ recipient cells and donor-derived Tact:** CTV-labeled *OT1C2GnT1^null^* donor cells were transferred into *PSGL1^null^* recipients with Ova antigen. On day 3 both recipient-derived CD8^+^ T cells and responding (CTV diluted), donor-derived, CD8^+^ Tact were evaluated in triplicate staining with biotinylated Meca79 (red *line)* or isotype-control *(blue line)* antibodies or no biotinylated 1^st^ antibody *(black line)*, either before *(left panel)*, or after applying EDTA-wash conditions used to strip PSL2 *(EDTA, right panel)*.

### PSL2 on Tact is docked on L-selectin

Since there is some overlap in PSel and LSel in recognition of PSelL and LSelL, the cell-extrinsic source of PSL2 and the preferential detection of PSL2 on Tact developing in PerLN vs spleen suggested that PSL2 might be a soluble ligand of LSel produced in PerLN. With this background, a primary candidate for PSL2 became Glycam-1, one of several secreted LSelL produced by HEV *(21, 22)* and that can bind PSel when immobilized on a surface (23). The *Glycam-1^null^* mouse was previously generated but abandoned when its physiological role as an LSelL could not be confirmed. We therefore re-derived and tested *Glycam-1^null^* mice for deficiency in loading PSL2 on Tact but observed no defect thus excluding Glycam1 as a candidate for PSL2 *(Supplemental Figure 1)*.

However, in the course of experiments to assess Glycam 1 as a PSL2 candidate, several observations indicated that LSel was indeed the dock on Tact for PSL2. *LSel*T cells access most PerLN poorly preventing direct comparison with *LSel^+/+^* cells but it has been reported that they can access mediastinal LN *(24)*. We found that entry of donor T cells into the tracheobronchial LN (TrBr LN)(25) was LSel independent and that these nodes were a relatively good source of donor Tact. Tracking dyes were used to distinguish donor *OT1 PSGL1’/’ LSel^+/+^* and *OT1 PSGLl^+/+^LSel^+/+^* Tact responding in the same TrBr LN of *PSGL1^nul1^* recipients and to assess the effect of LSel deficiency on PSL2 loading of Tact. As shown in *Figure 3B*, both *LSel^+/+^* and *LSel^-/-^* donor cells accessed the TrBr LN in equal numbers but PSL2 was not loaded significantly on *LSel^-/-^* Tact whereas *LSel^+/+^* Tact responding in these nodes cells loaded PSL2 as well as that seen in axillary lymph nodes.

LSel cell surface expression and LSel supported PerLN homing is gene dose dependent; *LSel^+/^-* T cells express half the LSel expressed by *LSel^+/+^* T cells *(Figure 3C)* but can still access PerLN albeit less efficiently than their *LSel^+/+^* counterparts *(26, 27)*. Whether such a two-fold difference in LSel expression would influence levels of PSL2 loaded onto Tact was assessed. PSL2 loading onto donor *LSel^+/^* Tact was compared with concurrently activated *LSel^+/+^* Tact recovered from the same PerLNs *(Figure 3D)*. Our data show that PSL2 loading on *LSel^+/^-* Tact was reduced relative to *LSel^+/+^* Tact.

LSel can be cleaved from the cell surface after T cell activation and, if required for PSL2 docking, such loss of LSel would be incompatible with PSL2 detection. The status of LSel on Tact was therefore assessed to further characterize the relationship between LSel and PSL2. As shown in *Figure 3E*, a fraction of donor-derived Tact were LSel^low/neg^, presumably due to LSel shedding, while a second subset remained LSel^hi^; PSL2 was only detected on a portion of these LSel^hi^ cells. Thus, there is heterogeneity in PSL2 expression on LSel^high^ cells and LSel^neg^ Tact are devoid of PSL2 expression.

Reagents that detect LSelL might specifically bind to PSL2 on Tact. One such LSelL-specific reagent is the IgM mAb Meca-79(28). Meca79 identifies most LSelL as it binds 6-sulfo sLex on extended core 1 0-glycans(10) but not 6-sulfo-sLex LSelL on core 2 branched O-glycans or N-glycans (12). We therefore examined if Meca79 would bind Tact and indeed specific binding was detected *(Figure 3F)*. This Meca79 signal was lost if cells were treated with the same EDTA washing procedure used to effectively removed PSL2.

Finally, PSL2 association with Tact requires Ca^++^, consistent with the requirement for this cation in selectin binding their ligands via their lectin domain. On the basis of the above observations we conclude that LSel is the dock for PSL2 on Tact and that PSL2 is a ligand for both LSel and PSel that can be simultaneously engaged by both selectins.

### Selectin recognition of PSL2

Whether PSL2 could serve as a ligand for ESel or LSel was explored. As shown in *Supplemental Figure 2* PSL2 was bound to some degree by all selectins and binding was eliminated by EDTA pre-washing. PSel recognition of decorated PSGL1 is dependent on sialic acid on sLex. As shown in *Supplemental Figure 3*, either PSel recognition of PSL2 and/or PSL2 docking onto LSel were vulnerable to neuraminidase indicating that sialic acid was required for one, or both, of these interactions.

### PSL2 transfer from recipient LNC to donor Tact

The cellular source of PSL2 is unknown but likely HEV inasmuch as HEV synthesize LSelL and are present in PerLN but absent in spleen. In experiments using Tact that expressed different combinations of PSGL1 and PSL2 we observed that Tact devoid of PSL2 rapidly acquire PSL2 if co-pelleted with PSL2+ cells. This phenomenon was explored in more depth. As shown in *Figure 4A* Tact from *OT1C2GnT1^nul1^* donors responding in *PSGL1^nul1^* recipients expressed PSL2 whereas the same donor cells responding in *C2GnT1^nuI1^* recipients did not. However, when ex-vivo PerLNC suspensions from the two recipients were mixed and co-pelleted, Tact from *C2GnT1^nuI1^* recipients could be stained with PSel-hlgG. This was consistent with PSL2 transfer from PerLNC to Tact. We then assessed if the in vitro acquisition of PSel+ signal required LSel expression on Tact. Spleen-derived *LSel^-/-^* Tact were compared with spleen-derived *LSel^+/+^* Tact for PSel staining before or after mixing with PSL2-competant PerLNC. The results in *Figure 4B* demonstrated that LSel was required for PSL2^neg^ Tact to become stainable with PSelhIgG, reinforcing the view that transferred PSelL was PSL2 and that PSL2 can be acquired by LSel on PSL2^neg^ Tact though contact with PerLNC.

**Figure 4:**
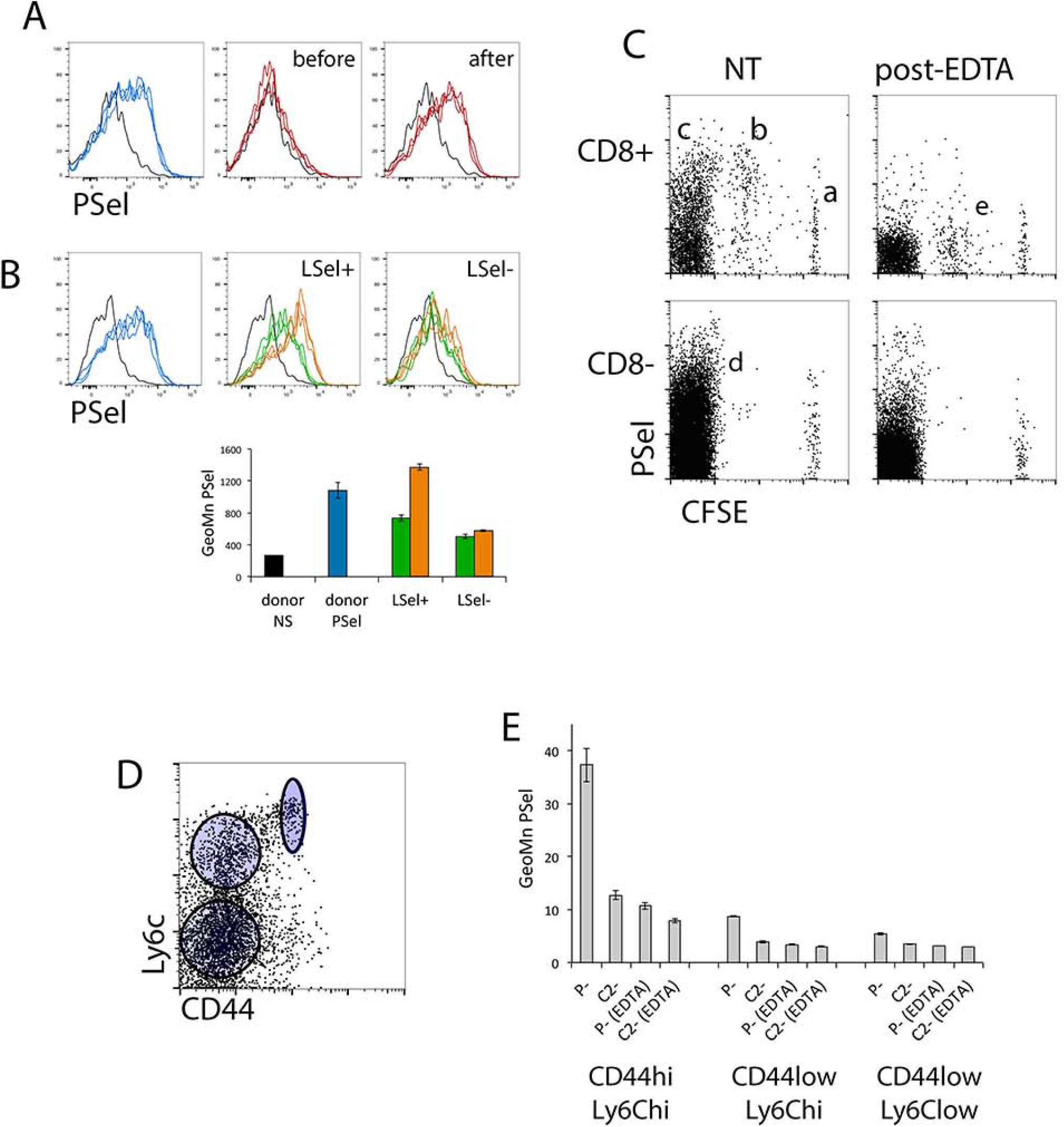
Tact can acquire PSL2 from PerLNC ex-vivo. **(A) Tact responding in C2GnT1^null^ recipients lack PSL2 and are not stained with PSel-hIgG - but become stainable after mixing with C2GnT1+ PerLNC:** *Left panel (blue)* CFSE-labeled *OT1C2GnT1^null^* Tact generated in PerLN of *PSGL1^null^* recipients stained in triplicate with PSel-hIgG to detect PSL2. No-PSel-hIgG staining controls *(black). Center panel (red)*, PSel-hIgG staining of Tact generated from the same donor cells labeled with CTV tracking dye and responding in PerLN of *C2GnT1^null^* recipients showing lack of PSL2 signal. *Right panel*, PSel-hIgG staining of Tact generated in *C2GnT1^null^* recipients (red), distinguished by CTV+ CFSE^neg^ fluorescence, after they were briefly mixed and 4x serially pelleted with whole LNC from the *OT1C2GnT1^null^ → PSGL1^null^* recipients whose donor-derived Tact were stained in the *left panel (blue)*. **(B) Acquisition of PSel stainability requires LSel expression on Tact:** PSel-hIgG stain of *OT1C2GnT1^null^*Tact generated in PerLN of *PSGL1^null^* recipients stained in triplicate with PSel-hIgG to detect PSL2, *left panel (blue line, donor PSel* in histogram below). No-PSel-hIgG staining controls *(black, donor NS in histogram below)*. PSel-hIgG stain of Tact obtained from spleen 3-days after either *OT1C2GnT1^null^ → C2GnT1^null^ (LSel+, center panel)* or *OT1PSGL^-/-^LSel^-/-^C2GnT1^null^* (LSel-, right panel) adoptive transfers, before *(green*) or after *(gold)* Tact were mixed and serially co-pelleted with whole PerLNC from the *OT1C2GnT1^null^ → PSGL1^null^* adoptive transfer shown in left panel. Geometric mean fluorescence of PSel-hIgG staining summarized in histogram. Standard deviation of triplicate stains shown. **(C) Recipient PerLNC also express an EDTA-removable signal detected by PSel-hIgG:** PSel-hIgG staining of CFSE-labeled *HY C2GnT1^null^* donor cells responding in male *PSGL^null^* recipients. CD8^+^ and CD8^-^ PerLNC were analyzed either directly *(NT)* or post-EDTA-wash *[post-EDTA)*. **(D) Memory phenotype CD8^+^ cells in ‘naïve’ PerLN:** Gated analysis of the CD4^neg^/CD19^neg^ (CD8^+^) subset of PerLN from naïve *PSGL1^null^* for expression of CD44 and Ly6c. **(E) Within the CD8^+^ subset of ‘naïve’ PerLN, only memory phenotype cells express PSL2:** PSel-hIgG staining of CD8^+^ PerLNC with gating as shown in (D) for CD44^high^Ly6c^high^, CD44^low^Ly6c^high^, and CD44^low^Ly6c^low^ subsets from naïve *PSGL1^null^ (P-)* or *C2GnT1^null^ (C2-)* mice, either before *(NT)* or after EDTA pre-washing *(EDTA)*. Standard deviation of triplicate stains shown.

### A PSL2 reservoir on PerLNC

Since PerLN cell suspensions collected by our methods would not likely contain HEV, the source of in vitro transferred PSelL shown in *Figure 4A* was unclear. *Figure 4C* illustrates PSel-hIgG staining of day-3 PerLNC after CFSE-labeled *HYC2GnT1^nuI1^* donor cells were adoptively transferred into male *PSGL^nuI1^* recipients. Nonresponding CD8^+^ donor cells failed to load PSL2 *(Figure 4Ca)* whereas responding donor Tact load with PSL2 *(Figure 4Cb)*. Importantly, PSel-hIgG staining was also detected on a subset of *recipient* CD8^+^ and CD8^neg^ cells in PerLNC *(Figure 4Cc/d)* and this PSel signal could be eliminated by EDTA pre-wash, paralleling the loss of PSL2 on similarly treated Tact *(Figure 4Cb/e)*, as well as the loss of Meca79 signals (LSelL) on both recipient CD8^+^ and donor-derived Tact after EDTA pre-wash shown in *Figure 3F*.

The phenotype of CD8^+^ cells binding PSel-hIgG from naïve PerLNC was explored. The CD8^+^ subset in PerLN was subdivided according to CD44/Ly6c phenotype *(Figure 4D)* and costained with PSel-hIgG. PSel-hIgG binding was highest on the memory Ly6C^high^ CD44^high^ subset of CD8^+^ T cells *(Figure 4E)*. As seen for PSL2 on Tact, binding was C2GnT1-dependent and eliminated by EDTA pre-washing. We have not conducted phenotyping of the CD8^neg^ PerLN subset staining with PSel-hIgG.

We conclude that the PSelL signal on CD8^+^ and/or CD8^neg^ PerLNC was the only plausible source of PSL2 ligand transferred in *Figure 4 panel A* since this PSelL signal was PSGL1-independent, EDTA-elutable, C2GnT1-dependent, and the only other detectable source of PSelL in the cell suspension available for in vitro loading on Tact. Although likely originating from HEV, the data suggest a ‘reservoir’ of PSL2 exists on cells in PerLN that can donate PSL2 to T cells undergoing activation.

### PSL2 and PSGL1 cooperate in Tact adhesion to platelets and to immobilized PSel-hIgG

Data presented above show that PSL2 is bound to LSel on Tact and that this complex could be recognized by PSel-hIgG chimera in solution. Physiological PSel is expressed on the luminal surface of activated endothelium and on activated platelets. We therefore assessed how effectively the PSL2/LSel complex would support Tact adhesion to PSel presented on activated platelets and how it would support Tact rolling/adhesion on immobilized PSel-hIgG. Moreover, it was unclear how PSL2 might cooperate with PSLG1 in these physical-adhesive processes.

Tact expressing PSGL1 and PSL2, PSL2 alone, or neither were generated in conjunction with EDTA pre-washing to resolve the contribution of PSL2 to platelet binding. Staining controls shown in *Figure 5A* illustrate how PSGL1 (EDTA pre-wash resistant) and PSL2 (EDTA pre-wash vulnerable) constitute the total PSelL expressed on Tact. The involvement of platelet PSel in platelet adhesion to Tact was resolved by comparing adhesion of *wt* platelets vs PSel deficient platelets isolated from *PSel^null^* mice. Data shown in *Figure 5B* indicated that PSL2 was able to support PSel-dependent adhesion between Tact and activated platelets and that PSL2 could cooperate with PSLG1 in additive manner for this physical interaction. In all cases, EDTA pre-washing of Tact reduced platelet binding mediated by PSel. There was very little background platelet binding when both PSGL1 and PSL2 were absent. We therefore conclude that PSL2 can provide adhesive support with PSGL1 for physical interaction between Tact and PSel on activated platelets.

**Figure 5.**
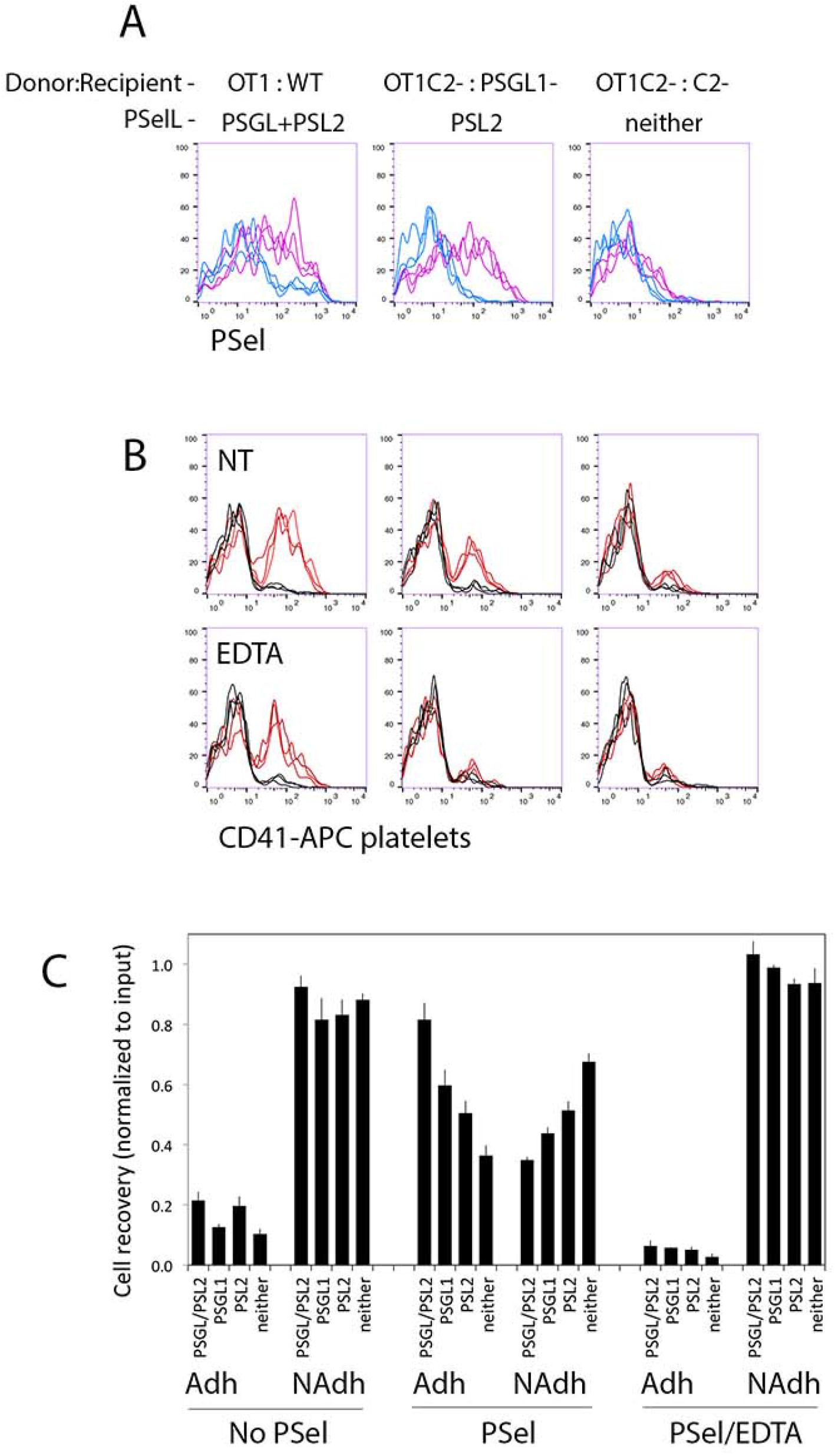
Adhesive function of PSL2. **(A) Platelet binding:** Donor/recipient combinations shown were used to generate Tact expressing PSGL1+PSL2, PSL2 alone, or neither, as shown. Staining with PSel-hIgG (PSel) was performed either without *(magenta trace)* or after *(blue trace)* EDTA pre-washing of Tact suspensions revealing the respective contributions of PSGL1 (EDTA resistant) and PSL2 (vulnerable to EDTA-stripping). Independent triplicate staining traces shown. **(B)** Platelets from *wild type (B6, red trace)* and *PSel^null^ (black trace)* mice were isolated, labeled with anti-CD41-APC, activated with thrombin, and fixed with paraformaldehyde. Platelets were then washed, mixed with untreated Tact *(NT)* or Tact previously washed with EDTA to remove PSL2 *(EDTA)*. Tact/platelet mixtures were then subjected to flow cytometry gating on viable Tact to assess acquisition of APC signal (platelet binding). Free platelets were excluded from analysis by forward/side scatter profile. **(C) Immobilized P-selectin mediates arrest of PSL2 expressing cells:** Responding (CTV-diluted) Tact from *OT1→ PSGL^null^, OT1→ C2GnT1^null^, OT1 C2GnT1^null^ → PSGL^null^, OT1 C2GnT1^null^ → C2GnT1^null^*, expressing PSGL1+PSL2, PSGL1, PSL2 or neither ligand respectively were compared for adhesion to immobilized PSel-hIgG chimera in V-bottom wells under shear generated by centrifugation. Wells were coated with blocking agent alone *(No PSel)* or with PSel-hIgG *(PSel)*. Cell aliquots were also centrifuged in wells pre-coated with PSel-hIgG but in the presence of EDTA to prevent selectin mediated binding *(PSel/EDTA)*. Non-adherent cells *(NAdh)* pelleting to the nadir were harvested directly and Tact adhering to the tapering well sides were subsequently harvested using EDTA (Adh). Tact in separately harvested non-adherent vs adherent fractions were quantified by FACS bead counting assay and normalized to input numbers of each Tact type. Triplicate counts of pooled quadruplicate samples were used to generate means and SD values shown.

The capacity of the PSL2/LSel complex to support Tact adhesion was also assessed by its ability to retard Tact rolling on a surface coated with PSel-hIgG. The method used to assess adherence entailed centrifuging Tact in V-bottom wells coated with PSel-hIgG, centrifugal g-force providing the shear against selectin engagement. Non-adherent cells pelleted to the nadir were harvested and then adherent cells retained on the tapered well walls were harvested separately after detachment with EDTA. Tact were quantified in both fractions by flow cytometry. Results shown in *(Figure 5C)* demonstrated that both PSGL1 and PSL2 contributed to Tact physical adherence in additive manner; there was no indication of adherence synergy between PSGL1 and PSL2 or a dominant contribution by either PSelL. Tact adhesion required imPSel and available divalent cation insofar as adhesion was prevented when EDTA was included during Tact pelleting. Results also suggested that a minor component of adherence to imPSel detected in this assay was independent of both PSGL1 and PSL2 insofar as Tact lacking both PSL2 and PSGL1 adhered above background as defined in wells lacking imPSel. We therefore conclude that PSL2 can provide adhesive support with PSGL1 for physical interaction between Tact and both activated platelets and immobilized PSel.

## Discussion

We describe the discovery and analysis of a PSGL1-independent PSelL that is acquired by responding CD8^+^ T cells after their activation in PerLN. This 2^nd^ PSelL, provisionally designated PSL2, contributed a significant proportion, variable but up to 50%, of the total PSelL expressed on primary activated CD8^+^ T cells observed with the models used here. PSL2 is an unusual SelL in that it is cell-extrinsic, contrasting with the canonical PSelL PSGL1 and other physiologically active cell-associated SelL whose production is cell-intrinsic. PSL2 expression on Tact was dependent on LSel and its docking onto Tact was Ca^++^ dependent, consistent with it binding LSel via its lectin domain. PSL2 is therefore a dual ligand for P-and L-selectins, able to physically bridge both selectins simultaneously. Importantly, PSL2 was the only PSelL other than PSGL1 detected on primary day-3 Tact in the models used.

PSL2 expression levels on Tact harvested from PerLN significantly exceeded that detected on Tact obtained from spleen. Given that LSelL are products of HEV and that PerLN have HEV while spleen does not, we anticipate that PSL2 is originally derived from PerLN HEV.

While the existence of additional selectin ligands has been anticipated (1) PSL2 has not been documented in prior analyses. The cell-extrinsic nature of PSL2, its absence on Tact generated in vitro, the rapid rate of its disappearance on cultured ex-vivo Tact, and the ease of PSL2 removal in media lacking Ca^++^ may account for why PSL2 has not been previously described. Furthermore, much of the efforts to identify SelL in the past were based on leukocytes (neutrophils and myeloid cell lines)(1) or in vitro expanded T cells that will not likely express PSL2. Our study focused on the primary CD8^+^ T cell response in mice but preliminary observations using the *OTII* mice suggested that some EDTA-strippable PSelL is present on CD4^+^ Tact.

The cell-extrinsic nature and the low availability of PSL2 have so far stymied efforts to identify it. As an extrinsic, likely HEV-sourced, LSelL PSL2 could be either secreted or cleaved from a membrane anchored LSelL. Multiple molecules have been identified as able to bind selectins, many of them with unresolved physiological roles as SelL. PSL2 may be one among those non-PSGL1 PSelL that have been previously identified *(TIM1, CD44, CD24, suIfatides, endogIycan, GaIectin1, GPIbα, heparan suIfate, nucIeoIin, thrombospondin)*. So far no soluble PSelL has been shown to be functional. PSL2 may alternatively be one among those non-PSGL1 LSelL that have been previously identified *(CD34, podocalyxin, endoglycan, MAdCAM-1, endomucin, nepmucin, nucleolin, VAP-1, CD44, perlecan, aggrecan, versican)*. Additional LSelL known to be secreted from HEV include *Glycam1, Sgp200, and Parm1*. As noted above, we re-derived the *GIycam1^null^* mouse but PSL2 detection on Tact generated in *GIycam1^null^* recipients was unaltered, ruling out Glycam1 as a PSL2 candidate. *Sgp200* and *Parm1* remain candidates for PSL2. *Sgp200* is a minimally characterized 200kD sulfated glycoprotein *(22, 29-31). PARM-1* is synthesized in HEV(32) and weakly secreted (33).

Twenty years ago criteria were proposed to resolve physiologically relevant selectin ligands (34). We summarize these criteria as follows, (i) that the ligand should be expressed at a time and place appropriate to a given selectin’s role in a process, (ii) selective removal, or absence, of the ligand from intact cells should render them unable to engage selectin in a biologically relevant interaction, and (iii) selectin should show specificity for the ligand and engage it with reasonably high affinity.

We considered PSL2 in the context of these criteria. PSL2 is expressed most prominently on primary Tact responding in PerLN where primary immune activation to infection is normally stimulated and drives formation of the canonical PSelL PSGL1. In our assay models PSL2 was expressed at significant levels, and in many cases comparable to PSGL1. PSL2 removal prevented PSel-dependent adhesion of Tact to immobilized PSel and to activated platelets. PSL2 binding to selectin was Ca^++^ dependent and of sufficient affinity/avidity to permit stable labeling with PSel-hIgG chimera and stable adhesion of cells to substrate bearing PSel. We conclude that according to these criteria PSL2 is likely to be physiologically relevant.

The unique properties that distinguish PSL2 among previously identified selectin ligands have generated a provocative landscape of insights and questions into how PSL2 may function and support selectin connectivity during recruitment. The logic of cell-extrinsic sourcing of a PSelL may seem paradoxical given conventional notions of SelL formation where T cells receive signaling input from stimulating dendritic cells that induce adhesion receptor expression.

What would be the utility of sticking a cell-extrinsic PSelL on LSel? Adding a second PSelL to the cell surface could simply enhance physical adhesiveness of Tact to substrates bearing PSel such as activated endothelia or activated platelets. Data presented here are consistent with PSL2 cooperating with PSGL1 for physical adhesion to PSel-bearing substrates. However, since Tact presents PSL2 in complex with cell surface LSel, LSel too will be indirectly engaged by PSel-bearing substrates. Given LSel’s signaling capacity and involvement in both homing and recruitment, it is reasonable to suspect that, aside from physical adhesion, additional consequences would accrue from LSel being drawn into engagement with1 PSel. Several known aspects of LSel distribution and function are salient to this discussion.

First, in several distinct systems LSel is limiting for important physiological outcomes including lymphocyte entry into lymph nodes*(8, 26, 27)*, neutrophil priming *(35)*, neutrophil recruitment in inflammation (36), neutrophil competence for response to chemotactic cues (37-39), and T cell mediated viral clearance (24).

Second, LSel is generally thought to exist as a monomer on the cell surface *(40, 41)*. However, there is evidence that upon T cell activation LSel may dimerize resulting in increased functional affinity for its ligands(42-46). For example, it is thought that to bind LSel physiologically, Glycam1 must either induce LSel clustering or bind to pre-clustered LSel(47). These observations were consistent with our observation that only responding donor Tact, but not quiescent donor cells, loaded PSL2 onto LSel.

Surprisingly, Tact acquisition of cell-extrinsic PSL2 occurring *in vivo* could be replicated *in vitro* by simply mixing and co-pelleting Tact generated in PSL2-’insufficienť *C2GnT1^nul1^* recipients with PerLNC from a PSL2-’sufficienť source eg. *PSGL1^nuH^*. Transfer of PSL2 occurred rapidly at 4°C and required LSel on receiving Tact. The only visible source of PSL2 for such in vitro transfer was a subset of both CD8^+^ and CD8^neg^ cells resident in PerLNC of our specific-pathogen-free mice (eg. *B6* or *PSGL1^nul1^)* that stained with PSel-hIgG. Among the CD8^+^ T cells in PerLN this PSL2 signal was present only on the memory CD44^high^ Ly6c^high^ subset. The effective acquisition of PSL2 by Tact from resident PerLNC suggested that the functional affinity of LSel for PSL2 was relatively high on Tact.

Third, PSel engagement of PSGL1 can stimulate integrin activation*(3, 48-51)* while LSel signaling can enhance responsiveness to chemokines (37-39). A functional complex between LSel and PSGL1 has been described in mouse(52) and human(53) neutrophils that is reportedly essential for PSel-stimulated integrin activation. Co-signaling though both PSGL1 and LSel in such a preformed complex might impact adherence and post-adherence Tact behavior during recruitment.

Fourth, it has been reported that O-glycan extensions on selectin ligands such as PSGL1 are required for their sorting to lipid rafts on leukocytes (54). Although present in both lipid raft domains and non-raft domains (55) PSGL1 signaling was dependent on lipid raft integrity suggesting that only raft-associated PSGL1 molecules are competent for signaling(49). At least in bone marrow leukocytes, LSel is excluded from raft domains (55) but perhaps, by ‘dressing’ LSel as a PSelL with PSL2 on Tact, LSel is ushered into signaling-competent raft domains and in proximity to PSGL1.

We therefore propose a model *(Figure 6)* where LSel is reconfigured during T cell activation enhancing its affinity/avidity for PSL2 extracted from the extracellular fluid phase (HEV sourced) or directly from contact with resident PerLNC. By acquiring PSL2, LSel carries the O-glycan signals for its relocation into raft domains in proximity to PSGL1. Colocalization of PSGL1 and LSel/PSL2 would present two ligands for engagement with PSel providing avidity and co-signaling enhancements ensuring that both integrin activation and chemokine responses are supported. The participation of PSL2 in support of LSel signaling needs to be verified but the downstream implications of cooperation between PSGL1 and the LSel/PSL2 complex could be significant. In the case of Tact where proper chemokine responsiveness is required for CD8 effector T cells locating infected cells in tissue*(56, 57)*, and antigen-encounter secures resident memory CD8^+^ T cell development(58), engagement of LSel in the primary response could contribute to the effective recruitment and further development of effector T cells.

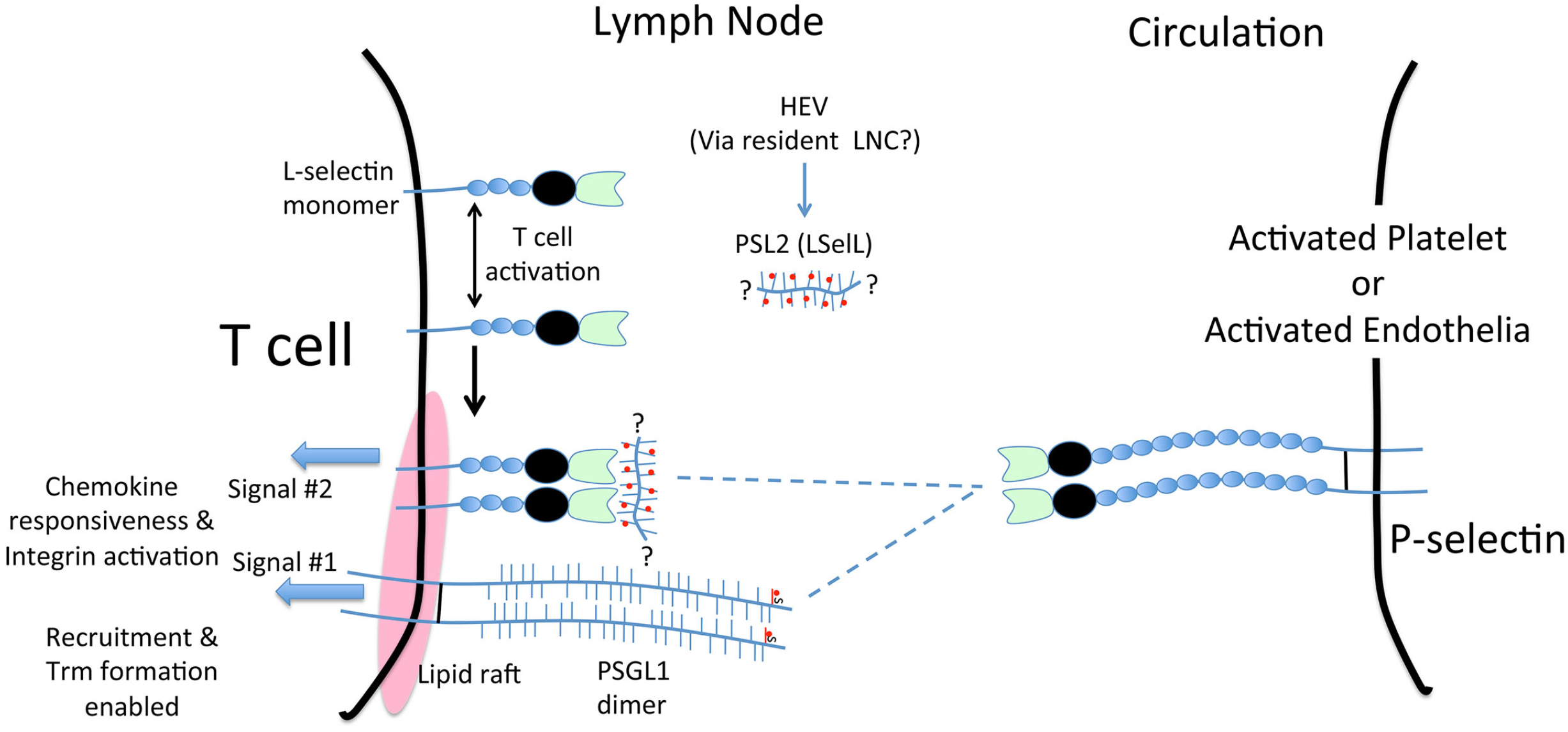

Observation of selectin-to-selectin connectivity is not without precedent. In contrast to mouse LSel, human LSel on both neutrophils and activated lymphocytes is directly modified by O-glycans for recognition by ESel*(59, 60)* enabling LSel:ESel engagement. Up to now this observation has been difficult to interpret as there was no prior precedent and little conceptual framework for why a selectin would itself be a selectin ligand, and why only in humans. The functional significance of ESelL on human LSel is still being explored. Importantly, in humans, ESel is thought to supplant the prominent role played by PSel in mice for tethering and recruitment(61). The murine Tact LSel:PSL2:PSel we describe may parallel the human LSel:ESel connectivity, both systems integrating LSel engagement upon leukocyte contact with endothelial selectins in inflammation. Analysis of PSL2 function offers a unique, technically accessible window into the functional importance of such LSel connectivity in vivo.

LSel was initially discovered in the context of T cell homing to lymph node via HEV, but analysis of *LSel^-/-^* mice also indicated a role for LSel during recruitment in experimental models of inflammation including DTH, allograft rejection, peritonitis, skin inflammation, and autoimmunity*(6, 51, 62-67)*. Understanding the nature and extent of LSel contribution to inflammatory cell recruitment has been complicated by (i) the deficit in naïve *LSel^-/-^* T cell homing to LN where T cells are stimulated, (ii) the absence of LSelL expression by endothelia in acute inflammation but inducible expression in chronic inflammatory settings*(2, 9, 11, 13, 68-70)*, (iii) variable LSel-supported recruitment through secondary capture *(1, 14, 66, 71)*, and (iv) LSel’s role in recruitment may also vary with inflammatory model and cell type under study.

It was recently reported in an influenza infection model that levels of LSel expression on T cells, early in the anti-flu response, correlated with both recruitment of CD8^+^ T effector (Teff) cells and influenza virus clearance(24). Mice with *null* or *shedding-resistant* mutations in LSel were susceptible/resistant respectively to flu infection relative to mice expressing wild type LSel. Teff priming, differentiation, and cytolytic function was intact while recruitment efficiency of CD8^+^ Teff and flu resistance was determined by levels of LSel. In a confounding twist, no LSelL (Meca79 binding) could be detected on vasculature of infected tissue. The prominence of L-selectin’s impact on flu resistance was unexpected; why were endothelial selectins insufficient for protection and how could LSel so influentially affect Teff recruitment without ligand expression on endothelia. The discovery of an LSel/PSL2 complex on Tact offers one possible explanation for a significant role of LSel in flu defense whereby PSL2 loading onto LSel may enable Teff recruitment through PSL2 tethering to endothelial PSel; the ‘missing LSelL’ on inflamed endothelium used by flu specific Teff could be PSel itself. Mohammed et al(24) also noted that Teff generated in vivo were recruited more efficiently than Teff generated in vitro, an observation consistent with involvement of PSL2. Based on our observations Teff generated in vitro would lack cell-extrinsic PSL2 and might shed LSel (PSL2 dock) more thoroughly rendering PSL2 loading of differentiated Teff inefficient after adoptive transfer.

Finally, we also demonstrated that PSL2 could support Tact adherence to activated platelets, the consequences of which deserve further investigation. Activated platelets express high densities of PSel(72) and represent a likely, physiologically potent, target of engagement for Tact bearing PSGL1 and LSel/PSL2. Such interactions might occur in circulation (73) and when either platelets or Tact are tethered to endothelia. Platelet-leukocyte interactions can promote recruitment *(74-77)* and support chemokinetic responses(78).

The identity of PSL2 remains unknown; its cell-extrinsic sourcing, low abundance and low frequency of Tact when gauged on preparative scales have stymied its identification. Although PSL2 is clearly capable of adhesive activity cooperating with PSGL1 for PSel-bearing substrates, the contribution of PSL2 during selectin-supported Tact recruitment in vivo will be confirmed with development of the PSL2 knockout mouse.

In summary, we present the discovery of a new PSelL, PSL2, acquired by Tact responding to antigen stimulation in PerLN. PSL2 is resolved as a cell-extrinsic P-selectin ligand bound to L-selectin on Tact. The LSel/PSL2 complex can engage PSel and co-operate with PSGL1 for adhesion to PSel-bearing substrates. The capacity to simultaneously bind and bridge PSel and LSel adds a new dimension to selectin connectivity and how, ‘dressed as a PSelL’, LSel may support recruitment. PSL2 will mediate signaling input driven by PSel and conducted through LSel on Tact. Our results also extend the scope of SelL function by demonstrating how the enigmatic cell-extrinsic soluble SelL can enhance connectivity among the selectins.

## Experimental Procedures

### Media and salt solutions

Routine media was designated I10 and included Iscove’s Modified Dulbeco’s Media (IMDM; Gibco Life Technologies #12440-046) supplemented with 10% fetal bovine serum (various suppliers), 100 U/ml penicillin, 100 U/ml streptomycin (Stem Cell Technologies), 2 mM glutamine (Sigma-Aldrich), and 5x10^−5^ M beta-mercaptoethanol (BioRad # 1610710). Dulbecco’s Modified Eagle Media (DMEM) Gibco #11965-084 supplemented with 20mM HEPES (Sigma Aldrich #H4034 pH 7.2) and 2 mg/ml BSA was used when staining with biotinylated antibodies. Calcium-free and magnesium-free phosphate buffered saline pH 7.4 (PBS) was prepared in-house. Hanks balanced salt solutions with Mg^++^ and Ca^++^ referred to here as ‘H+’ (Gibco Life Technologies #14025-092) or without Mg^++^ and Ca^++^ referred to here as ‘H-’ (Gibco Life Technologies #14170-112) were supplemented as indicated with bovine serum albumin (BSA; Sigma #A7906), EDTA pH 7.4, (ethylenediaminetetraacetic acid; BioRad #161-0728), CaCl2 (EMD #10035-04-8), MgCl2 (Fisher #M33-500), or MnCl2 (J. T. Baker Inc. #2540-04). H-B2 (H-with 2mg/ml BSA), H-B5 E2 (H-with 5 mg/ml BSA + 2mM EDTA), H-B2 E2 (H-with 2 mg/ml BSA + 2mM EDTA), H-B5 H10 Ca0.5 (H-with 5 mg/ml BSA + 10mM HEPES + 0.5 mM CaCl2).

### Mice

Mice aged 8-12 weeks were used for analyses. C57BL/6 (B6) mice were bred from founders obtained originally from Jackson Laboratory, Bar Harbour, Maine. *PSGL-1^null^* (B6.Cg-Selplgtm1Fur/J stock number: 004201) and *P-SeIectin^null^* mice *(PSeI^null^)* on the B6 background, and *Thyl.1* mice were also obtained from Jackson Laboratory. *C2GnT1^null^* mice(17) backcrossed with B6 mice beyond F7 were provided by Dr. Jamey Marth, Howard Hughes Medical Institute, University of California at San Diego, La Jolla, California. T cell receptor transgenic *OT1(79)* and *HY(80)* mice were backcrossed beyond F8 on B6 background. *LSel^nuĲ^* mice were provided by Dr. Steve Rosen (University of California at San Francisco). Mice were bred at the specific pathogen-free animal facility at the Biomedical Research Centre, University of British Columbia. Procedures employed in this study were approved by the Animal Care Committee at the University of British Columbia.

### In-vitro T cell stimulation

Dendritic cells from B6 or PSGL-1^nuI^ mice used for in vitro stimulations were prepared by differential adherence to plastic as previously described(81) except that spleen cell suspensions were not subjected to red cell lysis or filtered prior to plating. Dendritic cell-enriched suspensions were harvested after overnight detachment from petri plastic and either pulsed for 30 minutes at 37°C with 5 mg/ml ovalbumin in I10 or incubated for 60 minutes on ice with 10 μg/ml HY peptide KCSRNRQYL(82). Dendritic cells were then washed, counted, and cultured at 2x10^4^ cells/well together with either 3-10x10^4^ CD4-depleted HY thymocytes or 2x10^4^ OT1 LNC. After three days, cultures were harvested and stained with PSel-hIgG chimera.

### Antibodies, Selectin chimera staining, tracking dyes and flow cytometry

Antibodies: CD62L clone Mel-14 biotin (in house); anti-LSelL Meca79 (Biolegend #120804) and IgM isotype control (Biolegend #400803); CD44-FITC clone IM7.8.1 (in-house); Ly6C-APC (eBioscience #17-5932); CD8 clone 53-6.7 conjugates (APC, eBioscience #17-0081); Alexafluor-649 (in house); biotin (in-house); APC-EFluor-780, eBioscience 47-0081-82; PE, eBioscience #12-0081-85; FITC (in-house); Thy1.2 (TIB 107) Fab fragment biotin (inhouse); CD4-biotin clone GK1.5 (in-house);CD19-biotin clone 1D3 (in-house); CD41-APC (Biolegend #133914); Streptavidin conjugates (APC, eBiosciences #17-4317-82; PE-Cy7, eBioscience #25-4317-82; PE, BD-Pharmingen #554061). Propidium iodide (PI) was used at 200 ng/ml to label dead cells.

Prior to selectin-hIgG chimeras staining, ex-vivo lymphocytes were usually depleted sIg+ cells. Lymphocytes were stained for 30 minutes on ice with hIgG1 chimeras of mouse P-selectin (BD Pharmingen 555294), mouse E-selectin (R&D systems 575-ES), or mouse L-selectin (R&D systems 576-LS) at 5 μg/ml in I10 media ± 10mM EDTA. Cells were washed twice and stained with R-phycoerythrin conjugated goat-anti-human IgG Fc (Jackson ImmunoResearch #109-115-098), washed, and usually co-stained with fluorochrome conjugated CD8 and PI.

Two tracking dyes were used follow generation of donor-derived Tact 3 days after adoptive transfer. Prior to transfer ex-vivo donor cells to be labeled with tracking dyes were pelleted from I10 media and re-suspended in 37°C H+ containing either 2μM CFDA-SE (CFSE, Invitrogen C1157) or 5μM Cell Trace Violet (CTV, Invitrogen C34557), incubated at room temperature for 5 minutes, pelleted, washed once in H+, and injected into recipients.

Flow analysis with FacsCalibur and LSRII (Becton Dickenson). FCS file analysis was conducted with FlowJo software. Dead cells, debris, and aggregates were excluded from analysis by gating of forward scatter and PI^neg^ signals. Doublet discrimination applied to exclude doublets in data collected with LSRII.

### Adoptive transfers, Tact harvest, and counting

For OT1-based adoptive transfer responses, donor cell suspensions were prepared from pooled PerLN and mesenteric lymph nodes, depleted of surface Ig+ cells with Dynabeads sheep-anti-mIgG (Invitrogen #11031), and labeled with tracking dyes prior to intravenous transfer of 5x10^6^ cells in H+ with 1mg ova antigen into recipients. For HY-based male antigen specific responses, CD4-depleted donor HY thymus tissue was used. Donor thymus cell suspensions were prepared in I10 and depleted of CD4+ cells using anti-CD4 GK1.5 pre-loaded onto Dynabeads Sheep-anti-Rat-IgG (Invitrogen #11035) and labeled with tracking dyes prior to intravenous injection of 5x10^6^ donor cells per recipient. Three days later PerLN (and in some cases mesenteric LN or spleen) were harvested, cell suspensions prepared in I10, and depleted of sIg+ cells (as above) to enrich for Tact prior to selectin staining. To assess Tact responses in trachea bronchial LN (TrBr LN), OT1-based *LSel^-/-^* and *LSel+/+* donor cells were labeled with distinct tracking dyes, and co-injected intravenously into recipients. At the time of donor cell iv injections, recipient mice also received an intraperitoneal injection of 30 μl of stock 1 μm polystyrene bead suspension (PolySciences #17154) that had been pre-coated for 20 minutes with 10mg/ml ovalbumin in PBS and washed in PBS. Where indicated, donor Tact cell yields were monitored at day 3 using a FACS-based counting assay using 10 μm CML latex beads (Invitrogen C37259) as previously described (83).

### EDTA for selectin stain controls, EDTA pre-wash (PLS2 stripping), and cation titration

Two distinct types of EDTA treatments were conducted. Inclusion of 10mM EDTA during selectin chimera staining was used as a control for selectin binding via its lectin domain. EDTA *pre-wash* was also performed to ‘strip’ PSL2 from Tact. Cells at ≤2x10^7^/ml in I10 were stripped of PSL2 by adding EDTA to 10mM, rested on ice for 5 minutes, vortexed gently for 5 seconds, pelleted, re-suspended in H-B5 E2, rested another 5 minutes on ice, re-vortexed gently for 5 seconds, pelleted, re-suspended in I10, and filtered though a 70μm nylon screen.

### PSL2 rebinding assay

CD8^+^ *HY-C2GnT1^null^* donor Tact recovered from PerLN of male *PSGL1^null^ Thy1.1* recipients on day 3 were enriched by +’ve selection with biotinylated anti-Thy1.2 Fab and anti-PSGL1 4RA10 Fab antibodies using an EasySep Biotin Selection Kit (Stem Cell Technologies #18556). Cells were briefly washed 1x with 1 ml of H-B2, and then resuspended at 10^7^/ml in H-B2 E2 for 15’ on ice with brief gentle vortexing at 5’ intervals. Cells were then pelleted out of suspension and separated from the supernate (SUP) by centrifugation. The SUP was then spun at 16,000g for 1 minute to clear residual debris and supplemented with CaCl2 to 3mM Ca^++^ (yielding 1mM free Ca^++^). The aliquots of stripped cells were re-suspended in 75μl I10 media and divided into three equal aliquots incubated for 60’ on ice with (i) Ca^++^ replete *‘SUP’*, (ii) an equal volume of I10 media *(post-EDTA)*, or (iii) an equal volume of *MOCK SUP* generated as SUP but never exposed to cells. Cells where then washed in I10 and stained with PSel-hIgG.

### In vitro PSL2 transfer

*OT1C2GnT1^null^* donor cells were labeled with CFSE or CTV tracking dyes and adoptively transferred into either *PSGL1^null^* or *C2GnT1^null^* recipients to yield recipient PerLNC at day 3 containing CFSE-diluted PSL2+ Tact or CTV-diluted PSL2^neg^ Tact respectively. The latter cells were enriched for PSL2^neg^ donor Tact by depleting PerLNC with Dynabeads Sheep-anti-Rat-IgG (above) pre-loaded with rat antibodies specific for CD4 (GK1.5) and CD19 (ID3). PSL2 transfer to PSL2^neg^ enriched cells was observed after mixing and 4x repeated pelleting and re-suspension of 0.5x10^6^ of these with 10^6^ whole PerLNC from the former *OT1C2GnT1*^null^→*PSGL1^null^* adoptive transfer. Whether or not PSL2 transfer to PSL2neg Tact required using PerLN from adoptive transfer recipients versus from unmanipulated mice was unresolved at the time of manuscript submission.

### Platelet preparation

Platelets were prepared with all solutions at room temperature. Blood from B6 and PSel^null^ donors was harvested by heart puncture after Avertin anesthesia as follows. Approximately 1 ml of blood was drawn into a 3 ml syringe containing 150μl acid citrate dextrose solution (ACD; 22g/L trisodium citrate 2H2O, 8 g/L Citric acid H2O; 24.5 g/L dextrose), mixed by inversion, supplemented with 200μl PBS containing 7mM EDTA, mixed, and contents transferred into a 1.2 ml polypropylene cluster tube (#4401, Corning Incorporated, New York). Tubes were spun at 340g for 4 minutes and decelerated without brake. Platelets concentrated in the middle, cloudy, RBC-free layer (platelet rich plasma = PRP) were harvested in ≤ 200μl with a P200 Gilson pipette and added to 2 ml of PBS containing 7mM EDTA in a 5 ml polystyrene Falcon tube (Corning #352054), and respun at 1800g for 5 minutes with deceleration at lowest brake setting ‘1’. Supernatant was discarded and platelets re-suspended in 500μl H-B5 E2, labeled with 3μl of anti-CD41-APC for 5 minutes, added 2 ml H-B2, and re-pelleted at 1800g for 5 minutes at lowest brake setting ‘1’. Supernatant was discarded and platelets re-suspended in 2ml PBS with 0.1mM CaCl2. Bovine thrombin (Sigma #T6634) was added to 1 Unit/ml from a thawed 100 Unit/ml stock solution and the suspension incubated at 37°C for 4 minutes. Two ml of room temperature PBS containing 2% paraformaldehyde (Electron Microscopy Sciences #15710) was added, mixed, and incubated for 10 minutes at room temperature, transferred to a BSA-blocked 5ml tube containing 200 mg/ml BSA in PBS to yield a final BSA concentration of 10 mg/ml, and pelleted for 8 minutes at 1800g. Fixed and labeled platelets were then resuspended in H-B5 H10 Ca0.5, counted, pelleted, and re-suspended to 5x10^8^/ml in the same media. Adequate re-suspension at this step, by increasingly vigorous pipetting if necessary, was monitored by microscopy and was important for both dissociation of platelet aggregates and assay performance.

### Platelet binding assay

Day 3 harvest of PerLNC for platelet binding assay: Three days after injection of CFSE labeled donor cells later PerLNC harvested and single cell suspensions prepared, depleted of sIg^+^ cells, split, one half held in I10 while the other half being stripped with EDTA (as described above), resuspend in I10 to 5x10^6^/ml, and placed on ice. Once platelets were prepared (see below) PerLNC were then pelleted and re-suspended to 5x10^7^/ml in I10. Platelets (20μl at 5x10^8^/ml) and cells (20μl at 5x10^7^/ml) were combined in a 5 ml tube and incubated for 10 minutes at room temperature with gentle mixing every several minutes after which 380μl of a 1:1 mixture of I10 and H-B5 H10 Ca0.5 with 0.2 μg/ml PI was added. P-selectin dependent platelet binding to responding donor cells was assessed by comparing the extent of APC signal (anti-CD41-APC labeled platelets) acquired by donor Tact exposed to *PSel^+/+^* vs *PSel^-/-^* platelets. Data was acquired with gating for responding donor cells (CFSE-diluted) by CFSE signal level, and by excluding unbound platelets, large aggregates, and dead cells by gating against low forward-scatter events (platelets), high forward-scatter events (aggregates), and FL3^bright^ events (dead cells) respectively.

### Immobilized P-selectin-hIgG adherence assay

The cell adhesion assay applied was based on that previously described *(84, 85)* and modified as follows. Goat-anti-human IgG (Southern Biotech, 2040-01) was diluted to 5 μg/ml in pH 8.5 carbonate buffer (10 mM Na2CO3 and 35 mM NaHCO3), dispensed in 75μl volumes into V-bottom wells of 96-well V-bottom polystyrene plates (Nunc 249662) and incubated for 2 hours at 37°C. Wells were washed 4x with blotto (5% skim milk powder in PBS with 0.01% sodium azide), incubated in blotto for 30 minutes at 37°C, and for 10 minutes at 37°C in BB (3% BSA in PBS). Wells were washed 2x with selectin binding buffer (SBB = 2 mg/ml BSA in H+) and then incubated with 75μl per well of SBB ± 1 μg/ml P-Sel-hIgG chimera for 1 hour at room temperature and washed 4x with SBB prior to cell addition. Donor-derived Tact in adhesion assays were distinguished by CTV tracking dye labeling done prior to adoptive transfer. PerLN containing donor derived Tact 3 days after adoptive transfer were prepared in I10, filtered through 70μm nylon mesh, and depleted of sIg+ cells with Dynabeads sheep antimouse IgG (Invitrogen 11031). Cells were re-suspended to 10^7^/ml and on ice in I10 until used for adhesion assay. Just prior to assay, cells were diluted to 4x10^5^/ml cells in SBB + 15% FCS ± 10mM EDTA, aliquoted into prepared wells at 100μl/well and incubated for 15 minutes at 37°C. Plates were then spun for 10 minutes at 80g at room temperature with slow acceleration and minimal braking on deceleration. Nadirs from quadruplicate wells for each well group were individually harvested in 50μl drawn with a single channel P200 pipette and pooled into BSA-coated U-bottom wells. To remnants in each V-bottom well, 10μl of 50mM EDTA in H-B2 was added and mixed by light vortexing, incubated on ice for 4’, re-vortexed lightly for 5 seconds, re-incubated on ice for 4’, re-vortexed lightly for 5 seconds, and pelleted at 420g for 2 minutes. The supernatant was flicked out and the cell pellet re-suspended in 150μl H-B5 E2, transferred to BSA-coated U-bottom wells, transitioned to I10, stained with CD8-APC, and washed. Cell pellets were re-suspended with 120μl I10 containing PI and 5x10^3^ 10μm CML latex beads (Invitrogen C37259) used for counting. Cell counting by flow analysis was performed as described above using the LSRII and acquiring both beads and donor Tact based on light scatter and CTV florescence in triplicate counts of 10^3^ beads per sample.

## Author contributions

Conceptualization D.A.C. & H.J.Z.; Methodology D.A.C., H.J.Z.; Validation D.A.C., H.J.Z.; Formal Analysis D.A.C.; Investigation D.A.C., M.C.T. (Figs 2B,C); Resources H.J.Z.; Data Curation D.A.C.; Writing original draft D.A.C.; Writing Review & Editing H.J.Z. & M.C.T. & D.A.C.; Visualization D.A.C.; Supervision H.J.Z., D.A.C.; Project Administration D.A.C., H.J.Z.; Funding Acquisition H.J.Z., D.A.C.

## Acknowledgments

The authors wish to acknowledge Dr. Jamie Marth for providing the *C2GnT1^null^* mice, Drs. Maki Ujiie, Steve Rosen, Ninan Abraham, and Abdalla Sheikh for helpful discussions, the British Columbia Children’s Hospital BioBank, the KOMP Repository at University of California Davis for Glycam1^+/−^ sperm (stock #Glycam1 aF3), and Nicole Hofs at the Genetic Modeling Centre at the British Columbia Cancer Agency for re-derivation of the Glycam1^+/−^ mouse. Funding for this research was provided by the Canadian Institutes of Health Research (C.I.H.R.). The authors declare no conflict of interest that would influence results or interpretation of data presented.

---

**Supplementary Figure 1:**
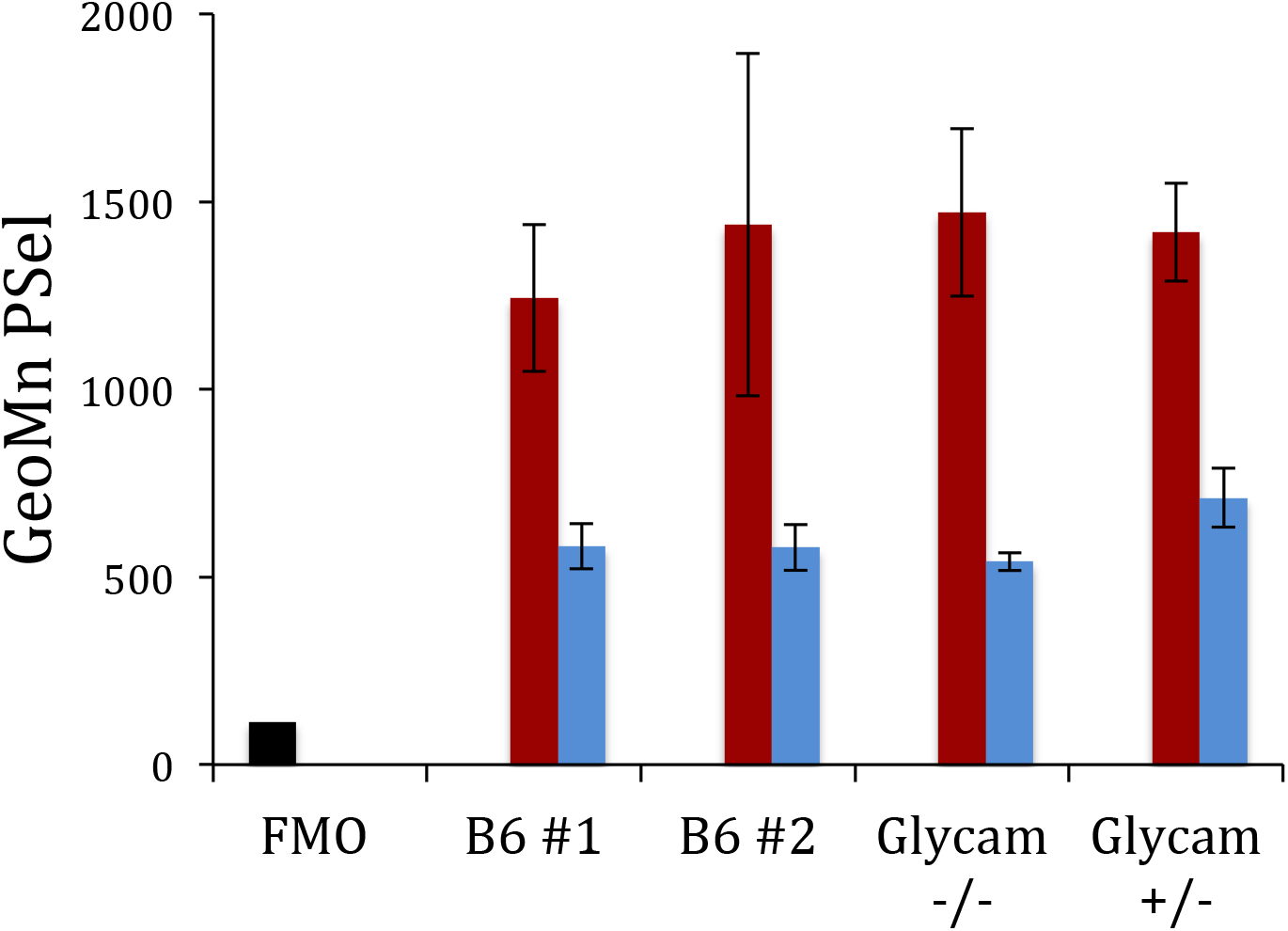
Legend: Tact responding in *Glycam*^-/-^ recipients load PSL2 normally. *OT1-C2GnTl*^-/-^ donor cells were activated for three days in two control B6 recipients, a *Glycaml*^-/-^ recipient, or a *Glycaml*^-/-^ littermate recipient. Peripheral lymph nodes were harvested and either untreated (red), or subjected to an EDTA-wash to strip PSL2 (*blue*), prior to staining with PSel-hIgG followed by anti-hlgG-PE and anti-CD8-APC. ‘Fluorescence-minus-one’ (FMO) control (*black*) staining of a B6 sample where only PSel-hIgG was omitted. Gated analysis of CD8^+^, pI^negative^ singlets shown. Figure shown is representative of three independent analyses. Error bars correspond to one standard deviation of triplicate stains of each sample.

**Supplementary Figure 2:**
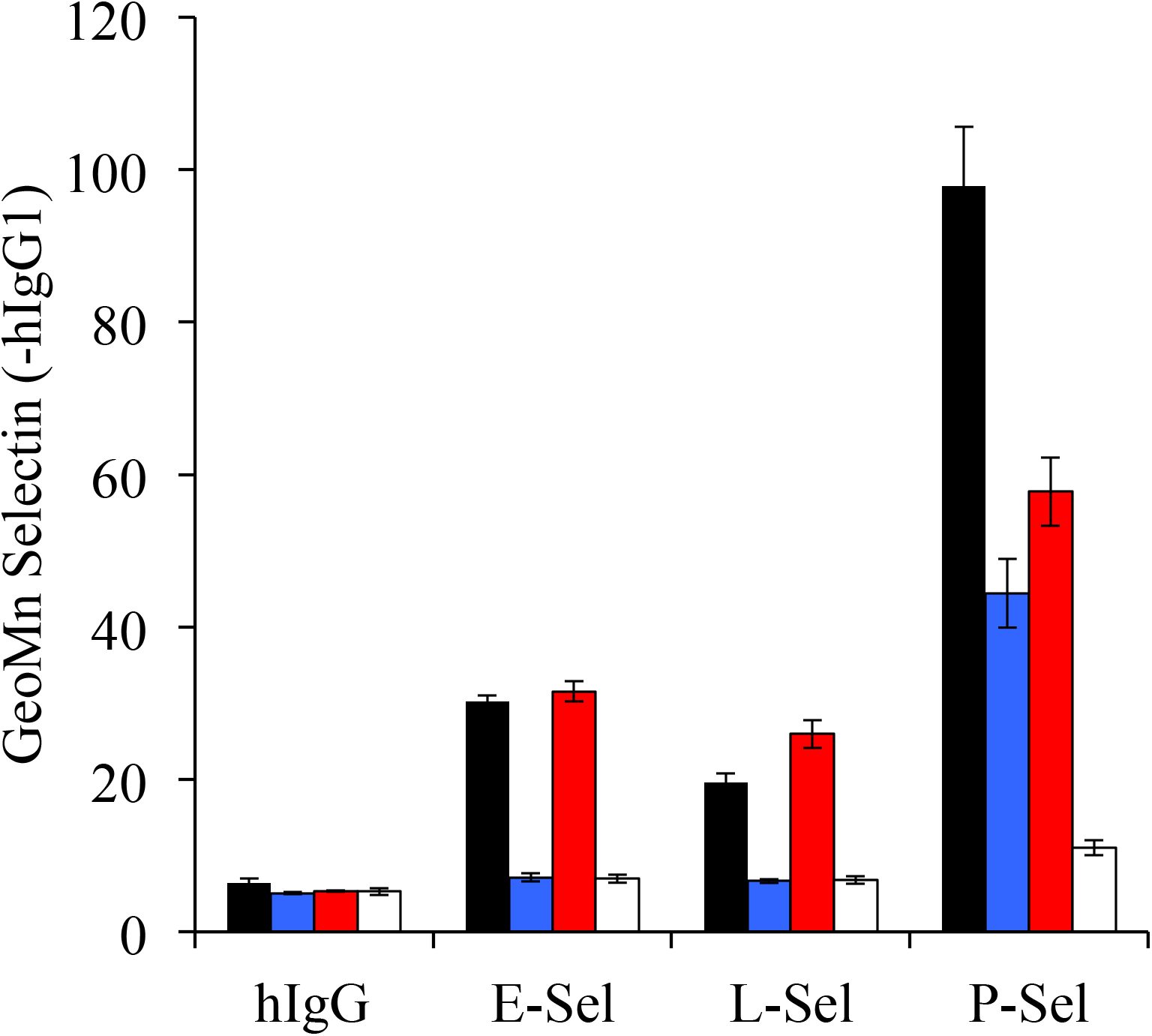
Legend: PSL2 is recognized by P, E, and L-selectin. CFSE-labeled donor *HY* or *HYC2GnT1^null^* cells were transferred into *PSGL1^null^ Thy 1.1* male recipients and recovered from PerLN on day 3. Aliquots of these cells were either left untreated or EDTA prewashed (PSL2-stripping conditions) and returned to Ca^2+^ replete I10 media for staining. In this way, four ex-vivo Tact preparations with distinct profiles of PSGL1 and PSL2 were compared for binding of of E-, L-, and P-selectin. *HY* donor (*black, PSGL1+PSL2)*, HY donor stripped *(blue, PSGL1 only), HYC2GnT1^null^* donor *(red, PSL2 only), HYC2GnTl^null^* donor stripped *(white, neither PSelL)*. Cells were and stained in parallel with unstripped counterparts using 5 μg/ml of hlgGl chimeras of each of the murine selectins followed by anti-hIgG-PE, CD8-APC and PI. PSL2 was bound to some degree by all selectins and binding eliminated by EDTA pre-washing. Selectin staining data shown as mean with standard deviation of three staining replicates of each population indicated gated on responding (CFSE-diluted), CD8^+^, PI^neg^ events.

**Supplementary Figure 3:**
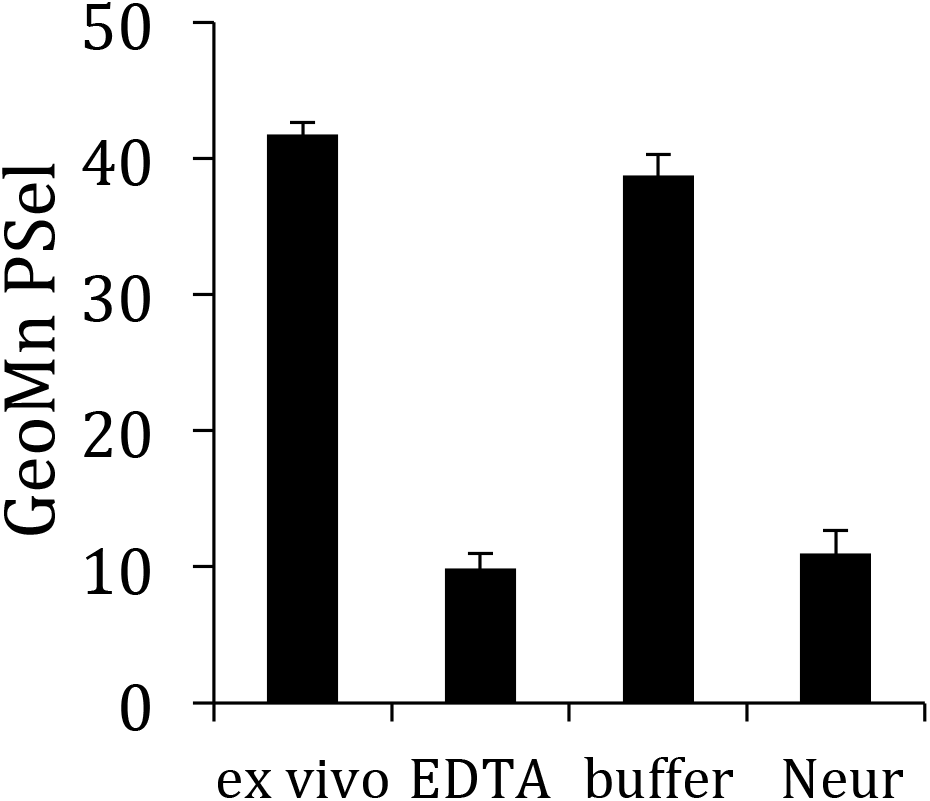
Legend: PSL2 detection on Tact by PSel-hIgG is dependent on sialic acid. *HYC2GnT1^nul1^* Tact were generated in *PSGL1^null^* recipients. PSel-hIgG staining of PSL2 on ex-vivo Tact *(ex vivo*) was prevented either by EDTA pre-wash *(EDTA)* or by exposure to neuraminidase *(Neur)* but not enzyme reaction buffer alone (*buffer*). Cells, 10^6^ cells per sample, were re-suspended in HC1 pre-acidified H^+^ pH 5.8 supplemented with 2.5% fetal calf serum. Neuraminidase Clostridium Perfringens (Roche #11 585 886 001 stock at 25 U/ml) was added to 125 mU/ml final concentration and the mixture incubated at room temperature for 15 minutes. Fetal calf serum was then added to a final concentration of 30% to reduce cell aggregation, and samples placed on ice for 5 minutes. They were then washed in Iı_0_ and stained with PSel-hIgG chimera.

